# Functional characterization of RNase H1 proteins in *Arabidopsis thaliana*

**DOI:** 10.1101/662080

**Authors:** Jan Kuciński, Aleksandra Kmera, M. Jordan Rowley, Pragya Khurana, Marcin Nowotny, Andrzej T. Wierzbicki

## Abstract

RNase H1 is an endonuclease specific towards RNA:DNA hybrids. Members of this protein family are present in most living organisms and are essential for removing RNA that base pairs with DNA. It prevents detrimental effects of RNA:DNA hybrids and is involved in several biological processes. We show that *Arabidopsis thaliana* contains four RNase H1-like proteins originating from two gene duplication events and alternative splicing. These proteins have the canonical RNase H1 activity, which requires at least four ribonucleotides for activity. Two of those proteins are nuclear, one is localized to mitochondria and one to plastids. While the nuclear RNases H1 are dispensable, the presence of at least one organellar RNase H1 is required for embryonic development. The plastid protein RNH1C affects plastid DNA copy number and sensitivity to hydroxyurea. This indicates that three genomes present in each plant cell are served by at least one specialized RNase H1 protein.

## INTRODUCTION

Double stranded nucleic acids, which contain deoxyribonucleotides on one strand and one or more ribonucleotides on the other strand are known as RNA:DNA hybrids. They are common byproducts of replication, transcription and other processes. Ribonucleotides within RNA:DNA hybrids are specifically removed by a class of endonucleases known as RNases H^1^. RNases H2 are multi subunit complexes capable of removing even individual ribonucleotides incorporated in double stranded DNA and have been studied in various eukaryotes, including plants^2,3^. RNases H1 act as monomers^4^ and require at least four ribonucleotide incorporated into double stranded DNA to bind the substrate^5^.

Among the substrates of RNases H1 are R-loops^6^ (RNA:DNA hybrid and a displaced ssDNA strand) which are often formed during transcription and replication^7–9^. These structures have also been implicated in DNA repair^10–12^, telomere maintenance^13^, IgG class-switch recombination^14^ and regulation of gene expression^9,15^. RNase H1 digests ribonucleotides within its substrate in a metal ion-dependent manner^16^, leading to single stranded DNA formation. Main feature of the proteins encoded by RNase H1 genes is the presence of the catalytic domain^16,17^. Additionally, some bacterial and most eukaryotic RNase H1 proteins contain a RNA:DNA hybrid binding domain (HBD)^18^.

Genes encoding RNase H1 proteins are present in the vast majority of living organisms, including Archea^19,20^, Bacteria^21^ and all kingdoms of Eukarya^1^. They are not essential in prokaryotes and lower eukaryotes but are required for survival in higher eukaryotes^1^. Eukaryotic genomes usually contain unique RNase H1 genes^1,21^, which may however be subject to alternative splicing^22^. *Arabidopsis thaliana* RNases H1 are encoded by three different genes with different predicted subcellular localizations^23^.

Among the three RNase H1 proteins in *Arabidopsis thaliana*, only the chloroplast-localized paralog has been studied so far^23^. It is essential for proper plastid development by maintaining the integrity of chloroplast DNA. It works with its interacting partner, DNA gyrase, to resolve transcription-replication conflicts and prevent DNA damage^23^. The role of the remaining two proteins remains unknown beyond the presumption that they resolve R-loops, which are relatively common in the *Arabidopsis* genome^24^.

Here, we characterize all RNase H1 proteins detectable in the *Arabidopsis* genome. We identify two ancient gene duplication events, which led to the formation of RNase H1 proteins targeted to various cellular compartments in monocots and dicots. A more recent duplication and alternative splicing produced four RNase H1 proteins in *Arabidopsis thaliana*, two of which are targeted to the nucleus, one to the chloroplast and one to the mitochondria. The proteins localized to endosymbiotic organelles are required for proper embryonic development. These proteins have the canonical RNase H1 activity and are involved in nucleic acid metabolism.

## RESULTS

### Origin of angiosperm RNases H1

Plant genomes have been shown to encode multiple RNase H1-proteins with different subcellular localizations^23^. To determine their evolutionary origins, we performed an in-depth phylogenetic analysis of plant RNase H1-like proteins. We first used BLAST to identify all plant proteins, which display sequence similarity to *Arabidopsis thaliana* AtRNH1C^23^ and contain both RNase H1 and RNA:DNA hybrid binding (HBD) domains. The list of identified proteins is in the Supplemental File 1. The identified proteins were subject to a simultaneous Bayesian alignment and phylogenetic analysis implemented using BAli-Phy package in order to integrate over the uncertainty in both the phylogeny and alignment^25^. In parallel, we predicted the subcellular localization of each identified protein using the TargetP prediction tool^26^. RNase H1-like proteins from angiosperms grouped into four distinct clades (Fig. 1, Fig. S1). The first group included proteins from monocots with mostly nuclear predicted localization. The second group included proteins from monocots with mostly chloroplast or mitochondrial predicted localization (referred to as organellar). The third group included proteins from dicots with mostly organellar localization. Finally, the fourth group included proteins from dicots with nuclear localization (Fig 1, Fig. S1, Fig. 2A). Phylogenetic relationships between these proteins indicate that the common ancestor of monocots and dicots had one RNase H1-like protein. Two independent gene duplication events then led to both monocots and dicots acquiring at least two proteins with distinct subcellular localizations.

**Figure 1.**
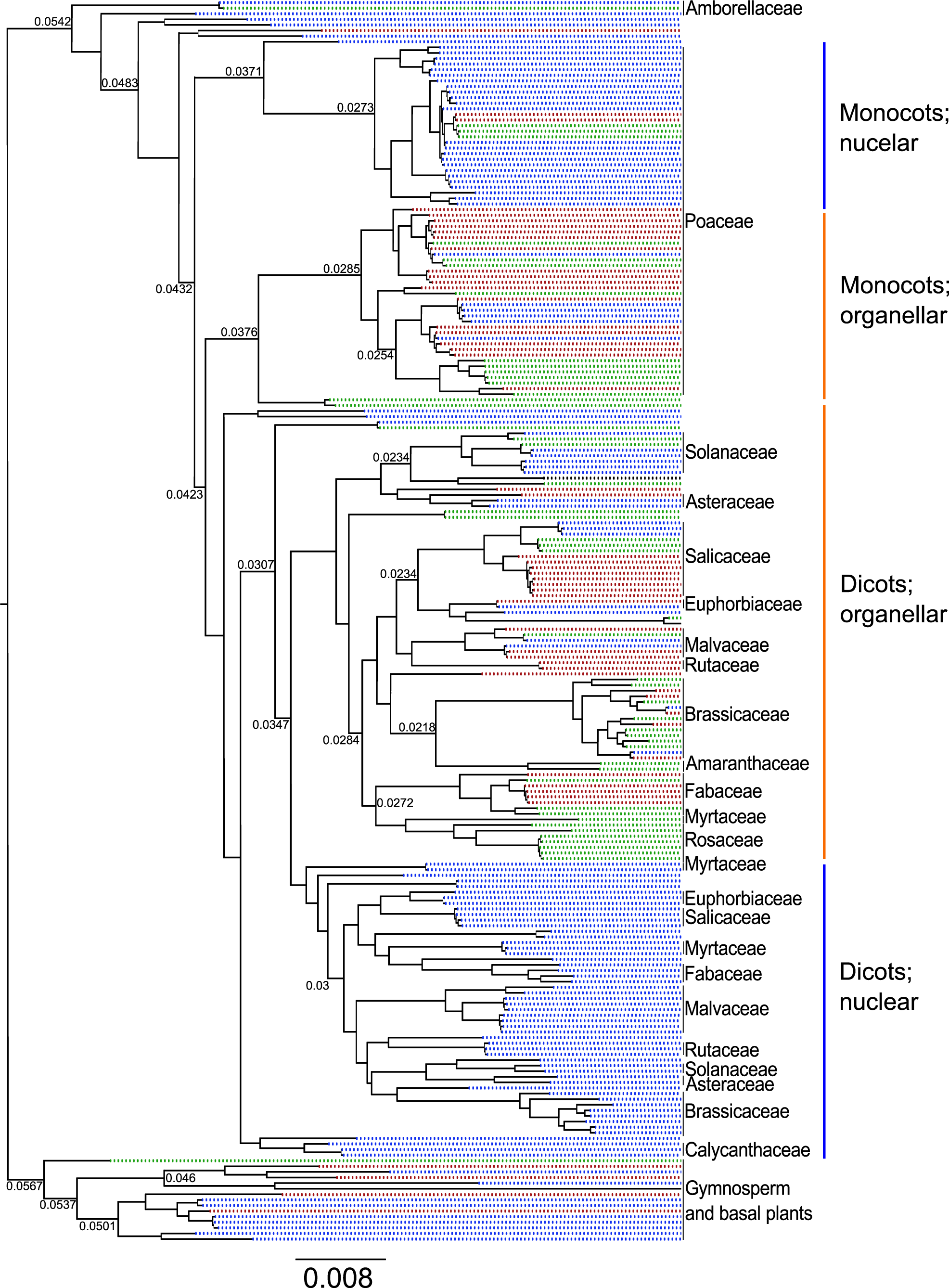
Evolutionary origin of plant RNase H1 proteins. Phylogenetic tree of all full length predicted RNase H1 proteins. Support values at tree branches are posterior probability scores, which integrate over the uncertainty in both the alignment and the phylogeny. Predicted protein localizations obtained using TargetP are marked with colors. Mitochondria – red, chloroplast – green, other – blue. A version of this phylogenetic tree with all species names and support values is shown in Fig. S1.

**Figure 2.**
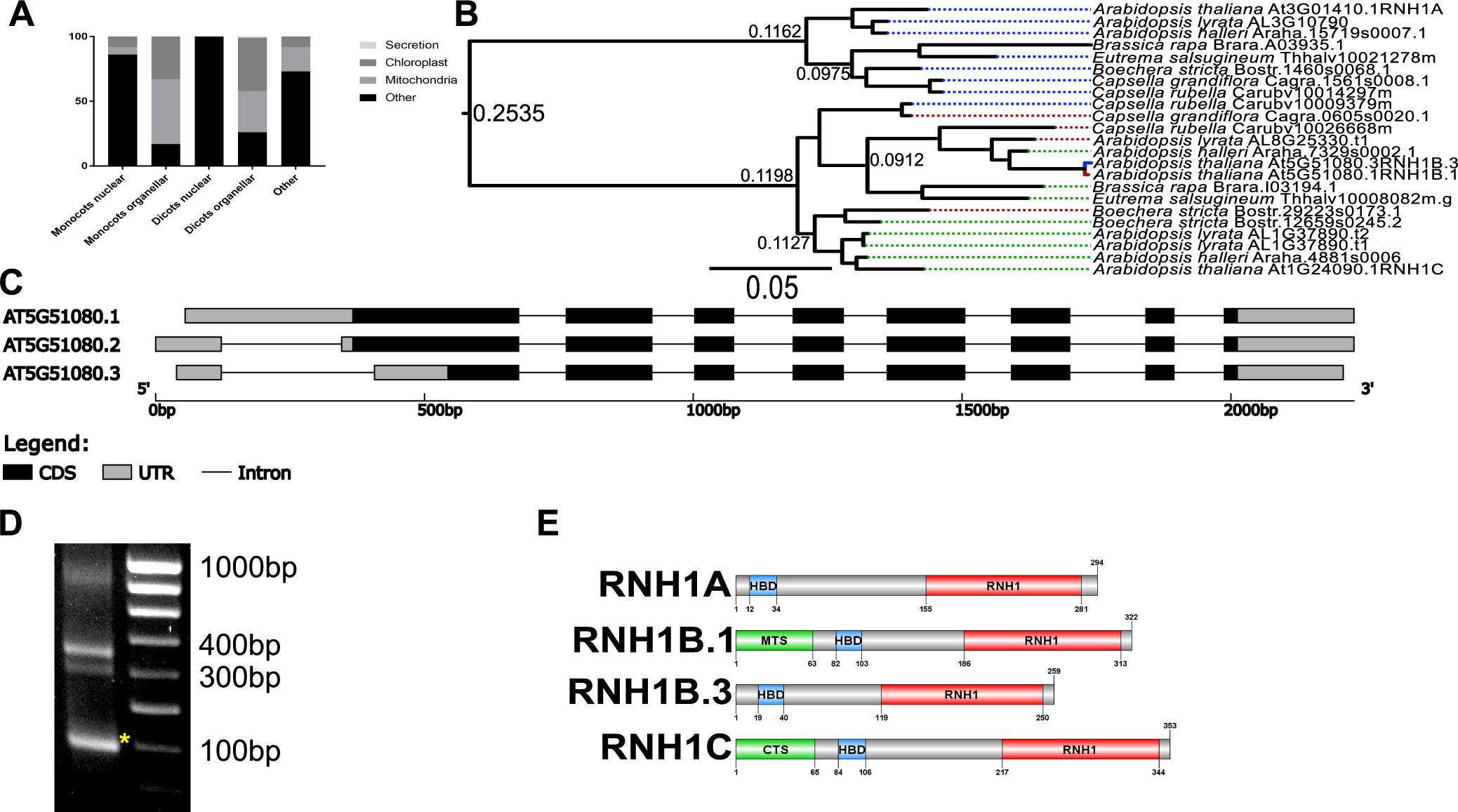
Origin of four RNase H1 proteins in *Arabidopsis thaliana*. **(A)** Predicted localization of RNase H1 proteins from distinct clades identified in Fig 1. **(B)** Detailed phylogenetic tree of all RNase H1 proteins in *Brassicaceae*. Support values and color-coded predicted localization like in Fig 1. **(C)** Splice variants of *RNH1B* predicted in Araport11. **(D)** 5’ RACE of *RNH1B*. Asterisk indicates a non-specific PCR product. **(E)** Domain composition of four RNase H1 proteins in *Arabidopsis thaliana*.

### Origin of four RNases H1 in *Arabidopsis*

*Arabidopsis thaliana* has been shown to contain three genes encoding RNase H1-like proteins: RNH1A (AT3G01410), RNH1B (AT5G51080) and RNH1C (AT1G24090). Products of these genes localize to the nucleus, mitochondria and chloroplasts, respectively^23^. While the split into nuclear and organellar proteins occurred early in dicot evolution (Fig. 1), the origin of two organellar proteins remains unknown. To determine the relationship between RNH1B and RNH1C we analyzed the phylogeny of RNase H1-like proteins in *Brasicaceae* (Fig. 2B). The phylogenetic tree identified one clade of nuclear and two distinct clades of organellar RNase H1-like proteins (Fig. 2B). This indicates that a second diversification event occurred in the common ancestor of *Brasicaceae*, which led to the formation of two organellar proteins. While in *Arabidopsis thaliana* these proteins are localized to mitochondria and chloroplasts, these specific localizations cannot be conclusively predicted for other *Brasicaceae*.

Multiple subcellular localizations of proteins produced from unique RNase H1 genes in animals are commonly determined by alternative splicing^27^. Araport11 genome annotation^28^ suggest a similar mechanism in *Arabidopsis thaliana*, where three splice variants of RNH1B (AT5G51080) have been identified (Fig. 2C). We confirmed these annotations using 5’RACE (Fig. 2D). While one splice variant (AT5G51080.1 and AT5G51080.2) encodes a full-length protein, the second variant (AT5G51080.3) has a truncated mitochondrial presequence region and is predicted to localize to the nucleus (Fig. 2E). We conclude that *Arabidopsis thaliana* contains four RNase H1-like proteins originating from two independent gene duplication events and alternative splicing.

### RNase H1-like proteins from *Arabidopsis* have canonical RNase H1 activity

Although *Arabidopsis thaliana* RNase H1-like proteins have extensive sequence similarity with RNase H1 proteins, their exact enzymatic activity remains unknown. RNH1C has been shown to remove binding sites of S9.6 antibody from chloroplast DNA^23^. This however, does not conclusively show RNase H1 activity of this protein. To determine if RNH1A and RNH1B are indeed RNases H1, we incubated the recombinant proteins with oligonucleotide substrates. Double stranded DNA and RNA was not digested by the recombinant proteins (Fig. 3AB). The same was true for oligonucleotides containing one or two ribonucleotides, which differentiates RNase H1 from RNase H2. Oligonucleotides containing four ribonucleotides were however digested by both RNH1A and RNH1B (Fig. 3AB). We did not test RNH1C because we were unable to produce a soluble recombinant protein. These results indicate that RNH1A and RNH1B have the canonical RNase H1 activity. Because of the extensive sequence similarity, especially within the catalytic domain, RNH1C is likely to have the same enzymatic activity.

**Figure 3.**
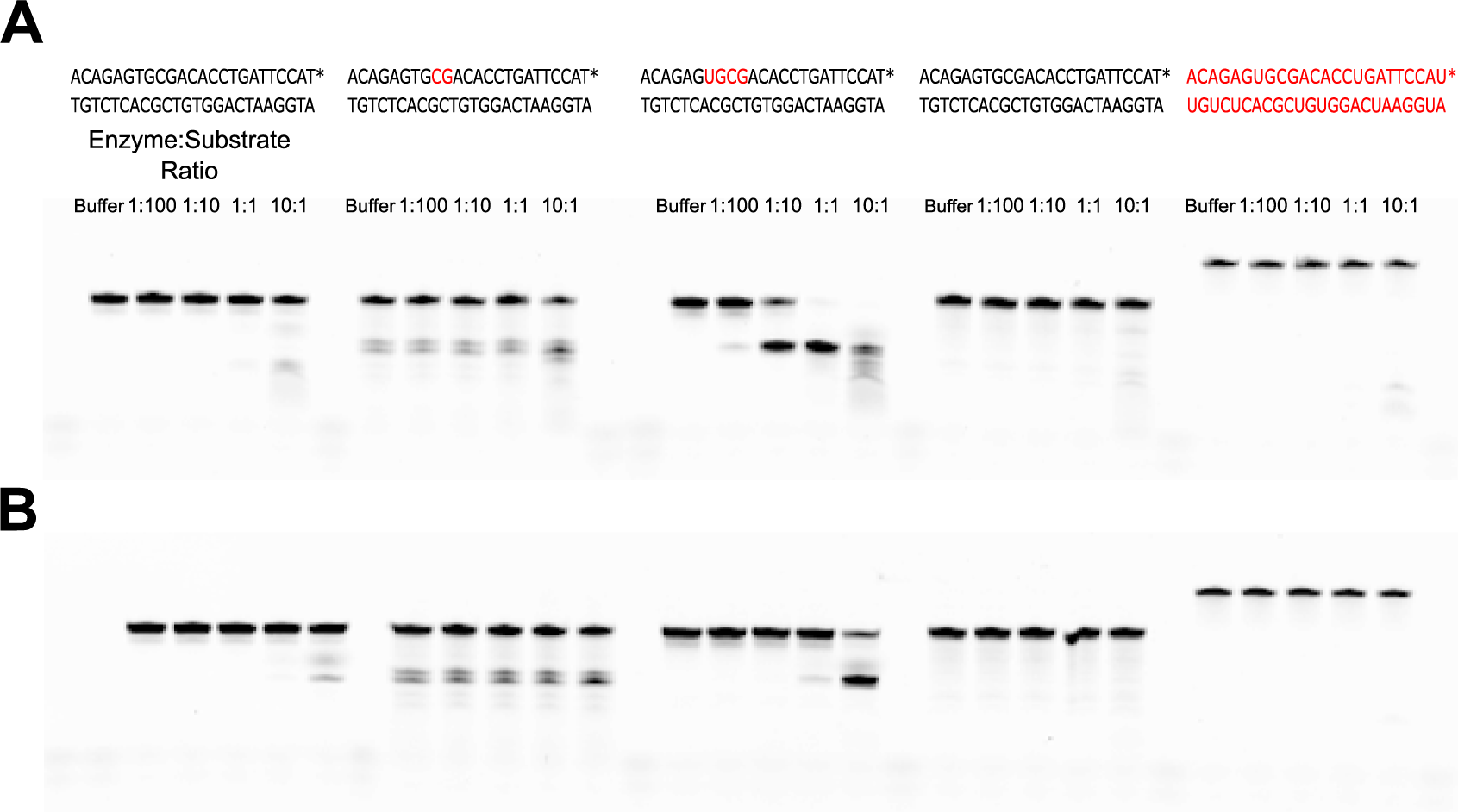
RNH1A and RNH1B proteins from *Arabidopsis thaliana* exhibit canonical RNase H1 activity *in vitro*. Recombinant RNH1A **(A)** and RNH1B **(B)** were incubated with labeled oligonucleotide substrates containing different combinations of deoxyribonucleotides (black) and ribonucleotides (red).

### RNH1A and RNH1B do not affect development

RNH1C has been shown to be required for proper chloroplast development^23^. To determine the roles of all three RNase H1 encoding genes, we obtained T-DNA mutants in RNH1A (SALK_150285C) and RNH1B (SAIL_1174_C11) as well as the previously published RNH1C (SAIL_97_E11). Single mutants *atrnh1a* and *atrnh1b* did not exhibit any obvious developmental phenotypes (Fig. 4A-C). *atrnh1c* mutant had the expected pale leaf and dwarf phenotype (Fig. 4D). The pale green phenotype of *atrnh1c* single mutant was confirmed by chlorophyll content quantification (Fig. 4I).

**Figure 4.**
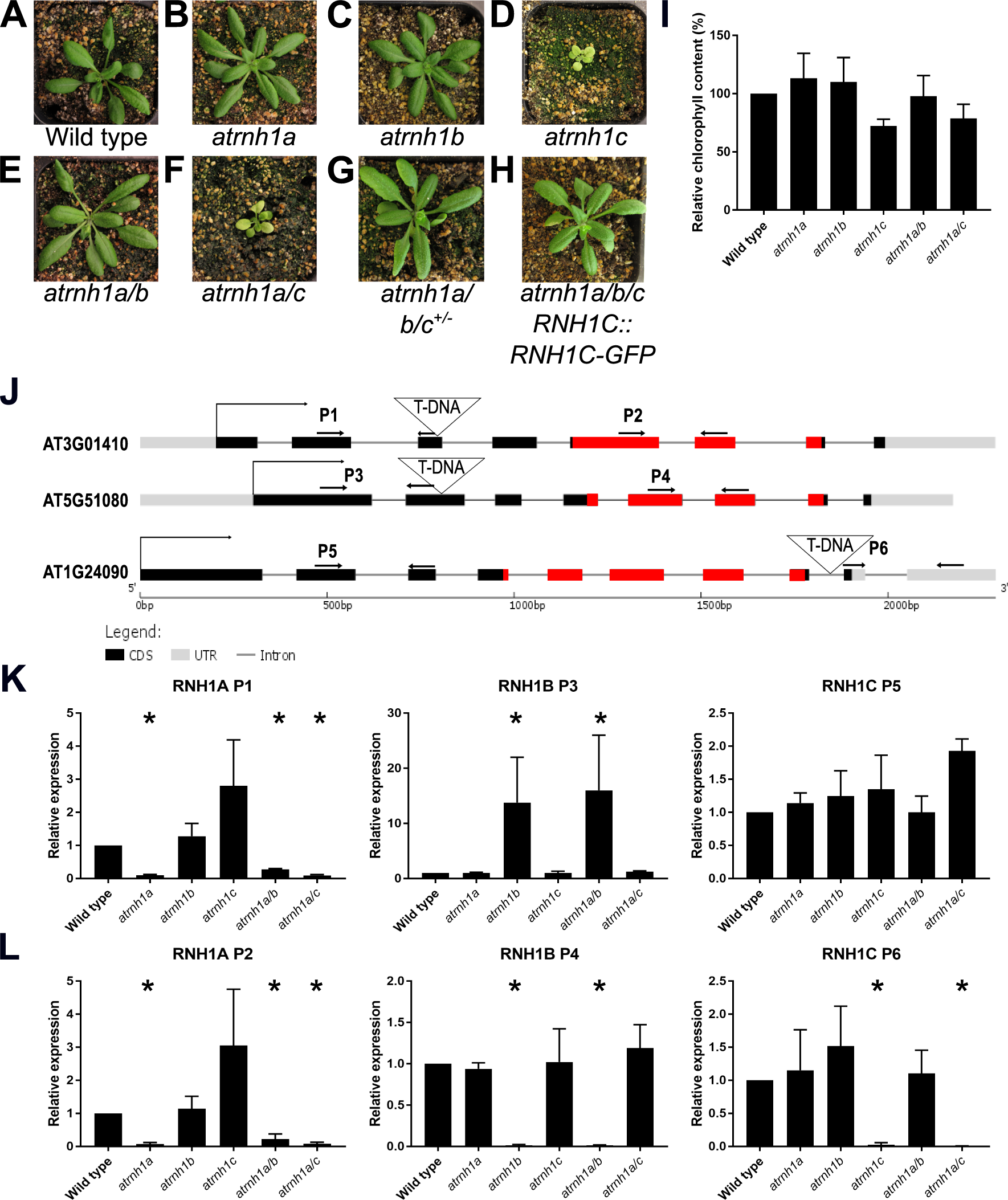
Effects of mutations in genes encoding RNase H1 proteins. Approximately 3-week-old plants of Col-0 wild type **(A)**, *atrnh1a* **(B)**, *atrnh1b* **(C)**, *atrnh1c* **(D)**, *atrnh1a, atrnh1b* double mutant **(E)**, *atrnh1a, atrnh1c* double mutant **(F)**, *atrnh1a, atrnh1b* double mutant heterozygous for *atrnh1c* **(G)** and *atrnh1a, atrnh1b, atrnh1c* triple mutant expressing RNH1C::RNH1C-GFP **(H). (I)** Relative chlorophyll content of Col-0 wild type and mutants in genes encoding RNase H1 proteins. Error bars indicate standard deviation from three biological replicates. **(J)** Schematic representation of RNH1 genes in *A. thaliana*. Arrows indicate START codons, triangles indicate T-DNA insertion positions. Red boxes indicate positions of the conserved RNase domain. **(K, L)** Expression of genes encoding RNases H1 measured **(K)** upstream of T-DNA insertion and **(L)** downstream of T-DNA insertion. Primer pairs are marked in (J). Error bars indicate standard deviation from three biological replicates. Asterisks indicate significant change in expression determined by 95% confidence intervals.

To determine if the studied T-DNA mutants may express truncated proteins, we performed RT-PCR with primers upstream or downstream of the T-DNA insertion sites (Fig. 4J). All mutants had strongly and significantly reduced RNA accumulation downstream of T-DNA insertions (Fig. 4KL). This indicates that these mutants are unlikely to produce truncated proteins containing the C-terminal RNase H1 domain. RNA accumulation upstream of T-DNA was reduced in *atrnh1a*, unchanged in *atrnh1c* and strongly increased in at*rnh1b* (Fig. 4KL). This indicates that a truncated N-terminal fragment is unlikely to be produced in *atrnh1a*. Truncated N-terminal fragments may be produced in *atrnh1b* and *atrnh1c*, however *atrnh1b* is not expected to contain full length RNase H1 domain. On the other hand, *atrnh1c* may produce a truncated protein including the RNase H domain (Fig. 4J), which indicates that the C-terminal part of the protein is important for its function. The strong increase of upstream RNA accumulation in *atrnh1b* may indicate the presence of an autoregulatory mechanism within *RNH1B*.

### Presence of both RNH1B and RNH1C is required for viability

Because RNases H1 are required for viability in animals^27^, we tested the phenotypes of all combinations of RNH1 double mutants. The *atrnh1a, atrnh1b* double mutant did not show any visible developmental phenotypes (Fig. 4E) and the phenotype of the *atrnh1a, atrnh1c* double mutant was similar to the *atrnh1c* single mutant. Phenotypic similarity of *atrnh1c* single mutant and *atrnh1a, atrnh1c* double mutant was confirmed by chlorophyll content quantification (Fig. 4I). Despite several attempts we were unable to obtain the *atrnh1b, atrnh1c* double mutant. Among 211 F1 plants obtained from crossing *atrnh1b*^*-/-*^ with *atrnh1c*^*+/-*^ we found no viable double homozygous mutant. However, the cross resulted in about 25% of seeds not developing properly (Fig. 5A-C), which likely are the double mutants. These improperly developing seeds contain aborted embryos which stop developing at approximately heart-shape stage of embryonic development (Fig. 5DE). These results suggest that while the nuclear RNH1A is not required for viability, the presence of at least one organellar protein RNH1B or RNH1C is essential.

**Figure 5.**
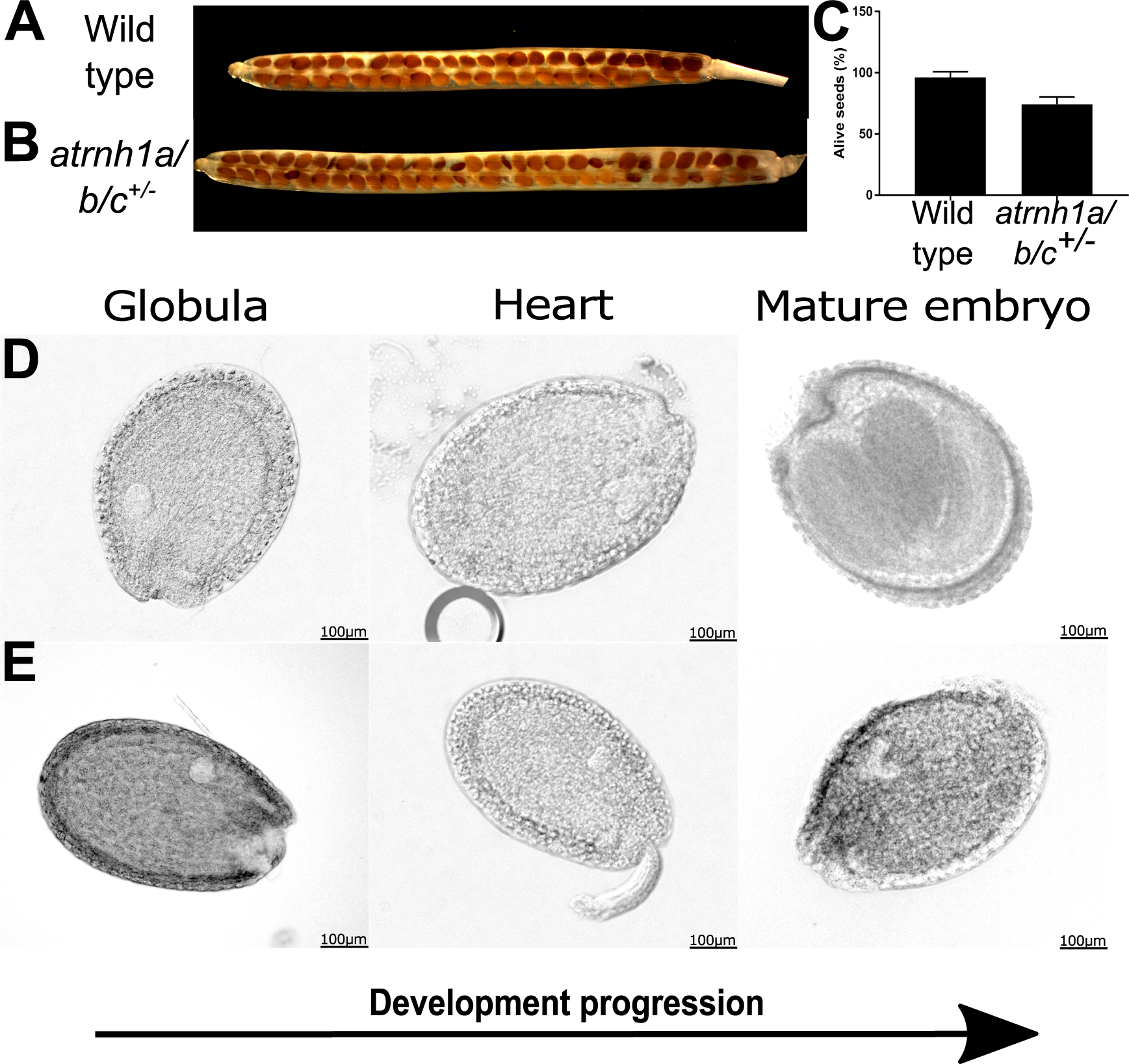
At least one organellar RNase H1 is required for embryo development. **(A)** Mature silique of Col-0 wild type. **(B)** *atrnh1a, atrnh1b* double mutant heterozygous for *atrnh1c* (*atrnh1a/b/c*^*+/-*^). **(C)** Percentage of properly developing seeds in siliques. Error bars indicate standard deviation from three biological replicates. **(D)** Development of a Col-0 wild-type embryo. **(E)** Development of an aborted embryo.

### RNH1C is involved in chloroplast nucleic acid metabolism

RNH1C has been shown to be involved in chloroplast DNA maintenance^23^. To test if RNases H1 expressed from the three *Arabidopsis* genes affect DNA copy number in various cellular compartments we performed real time PCR with primers specific to nuclear, mitochondrial and chloroplast genomes. *atrnh1c* mutant and *atrnh1a, atrnh1c* double mutant contained approximately three times more chloroplast DNA than wild type mutant plants (Fig. 6A-C). This result indicates that RNH1C protein is involved in controlling DNA copy number in chloroplasts, which is consistent with its postulated role in chloroplast genome maintenance^23^. Surprisingly, *atrnh1a, atrnh1b* mutants and the *atrnh1a, atrnh1b* double mutant did not affect the relative content of the three genomes (Fig. 6A-C). This indicates that nuclear and mitochondrial RNases H1 have different roles than the chloroplast RNase H1. Alternatively, some or all RNase H1 proteins may be targeted to more than one compartment.

**Figure 6.**
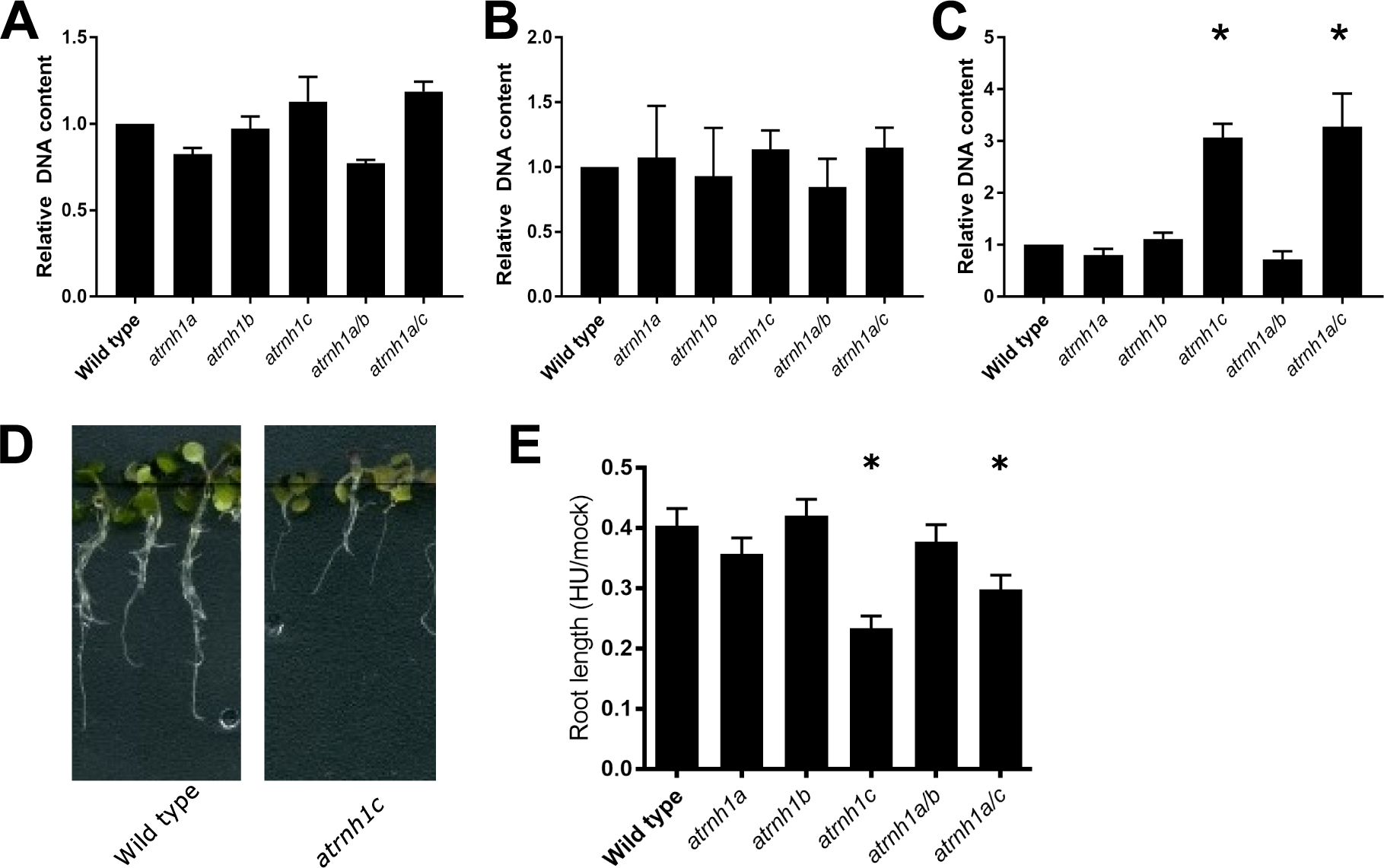
Loss of RNH1C leads to increased chloroplast DNA content and hypersensitivity to hydroxyurea. Relative levels of nuclear **(A)**, mitochondrial **(B)** and chloroplast **(C)** DNA in Col-0 wild type *atrnh1* mutants. Asterisks indicate significant changes with p<0.05. **(D)** Phenotype of plants subjected to HU stress. **(E)** Relative root length of Col-0 wild type and *atrnh1* mutants upon 2mM hydroxyurea treatment. Asterisks indicate p<0.0001determined using 2-way ANOVA.

The increased amount of chloroplast DNA in *atrnh1c* mutants might be a result of impaired DNA replication. To test this possibility, we treated wild type plants and RNase H1 mutants with replication stress and measured root length. To induce replication stress, we applied hydroxyurea^2,3,29^, which is known to inhibit replication by inhibiting rNTP reductase and therefore decreasing the amount of available dNTPs^30^. Treatment with 2mM hydroxyurea caused a decrease of root length in wild type plants to approximately 40% of root length in untreated plants (Fig. 6 E). *rnh1a* and *rnh1b* mutants had no effect on hydroxyurea sensitivity (Fig. 6 E). However, roots of *atrnh1c* mutant and *atrnh1a, atrnh1c* double mutant grew to less than 30% of root length in untreated plants (Fig. 6 E). This indicates that RNH1C affects sensitivity to hydroxyurea and is likely to be involved in DNA replication.

## DISCUSSION

RNase H1 proteins are known to digest the RNA component of RNA:DNA hybrids and this substrate must contain at least four sequential ribonucleotides^16^. In contrast, RNase H2 complexes only require a single ribonucleotide incorporated into double-stranded DNA for activity^1,31^. Therefore, RNase H2 has a broader substrate range and is partially redundant with RNase H1^32^. We demonstrate for the first time that *Arabidopsis thaliana* RNH1A and RNH1B have the canonical RNase H1 activity. Because the presence of RNase H2 complexes has been previously shown^2,3^, this indicates that plants, like other eukaryotes, have both RNase H1 and H2. Our results suggest that all three RNase H1 genes identified in the *Arabidopsis thaliana* genome encode canonical *bona fide* RNase H1 proteins.

*Arabidopsis thaliana* contains four RNase H1 proteins. Their evolutionary origin may be traced down to three events. The first event occurred in the common ancestor of dicots, where nuclear and organellar paralogs have been formed. An independent event in the common ancestor of monocots (or possibly the common ancestor of all angiosperms) led to the formation of similar, yet evolutionarily independent nuclear and organellar paralogs in monocots. Nuclear and organellar RNases H1 do not display any obvious structural differences beyond the presence or absence of a transit peptide/presequence. Therefore, it is unknown if they are functionally distinct beyond having different subcellular localizations and expression patterns.

The second event leading to the formation of four RNase H1 proteins in *Arabidopsis thaliana* occurred in the common ancestor of *Brassicaceae*. The ancestral organellar protein diversified into two proteins, which are represented by RNH1B and RNH1C in *Arabidopsis*. Although these proteins are localized to mitochondria and chloroplasts^23^, their ortologs in *Brassicaceae* do not have consistent predicted localizations and the functional impact of this event remains unknown. The third event is alternative splicing of RNH1B, which results in translation of proteins predicted to localize to mitochondria or the nucleus. This mechanism is reminiscent of dual localization of metazoan RHases H1^22^. Overall, our results are consistent with recent evidence that each genome within *Arabidopsis thaliana* cells has at least one RNase H1^23^. However, the nuclear genome is served by two RNase H1 proteins. Additionally, most eukaryotes, including *Arabidopsis thaliana* contain RNase H2 complexes, which are nuclear localized and at least partially redundant with RNases H1^2,31,32^. This may suggests that the nuclear genome requires the presence of several activities resolving RNA:DNA hybrids.

Our results indicate that at least one organellar RNase H1 is needed for proper embryonic development. This is consistent with data from metazoans, where RNase H1 is crucial for proper mitochondrial DNA replication during embryo development^27,33,34^. Viability of single *atrnh1b* and *atrnh1c* mutants may speculatively be explained by dual (mitochondrial and plastid) localization of those proteins, which is not an uncommon phenomenon^35,36^. Alternatively, defects in both mitochondria and chloroplasts may have a synergistic effect in embryonic development.

RNase H1 has been shown to be required for genome maintenance in chloroplasts^23^. Our observation that plastid DNA copy number is substantially increased in *atrnh1c* may indicate that genome instability in *atrnh1c* leads to overamplification of the entire genome. Because the plastid genome is copied by a combination of replication and recombination^37^, this overamplification may rely on homologous recombination. Interestingly, loss of mitochondrial RNH1B does not affect the level of mitochondrial DNA. This may suggest that, contrary to animals, plant mitochondria contain enzymes at least partially redundant with RNH1s.

The reported role of RNH1C in release of replication-transcription conflicts and chloroplast DNA integrity^23^ is consistent with our observations that *atrnh1c* mutation increases sensitivity to hydroxyurea. This may be interpreted as evidence of disrupted DNA replication in *atrnh1c*. This result should however be interpreted carefully because hydroxyurea inhibits synthesis of deoxyribonucleotides and increases misincorporation of ribonucleotides^29,30^, which may have a replication-independent effect on RNase H1-dependent processes. Additionally, hydroxyurea may cause oxidative stress ^38–40^, which could also have a replication-independent effect.

Our results raise questions about the roles of nuclear and mitochondrial RNases H1 in *Arabidopsis thaliana*. Single mutations in genes encoding these proteins do not result in visible phenotypes, which may likely be attributed to protein redundancy and localization to multiple cellular compartments. Resolving the roles of those proteins remain an important goal for future studies.

## MATERIALS AND METHODS

### Plant material and oligonucleotides

Plant lines used in this study: Col-0 (CS 70000), *atrnh1a* (SALK_150285C), *atrnh1b* (SAIL_1174_C11), *atrnh1c* (SAIL_97_E11) and crosses between abovementioned. Oligonucleotides used for expression analysis: P1: 5’-TTTAGTTTGGGTTGATGGGTTCC-3’ + 5’-CACACTCATTGCAGGATGTGATAC-3’; P2: 5’-ATCTCTTAGACGGGGAAGATTTGT-3’ + 5’-AATGCAAGTCATGTCAAAGATGAT-3’; P3: 5’-TCAATAATGCAAGTTTCATATGAGGT-3’ + 5’-GTTTGGAGCTCTTACACCTTGTCT-3’; P4: 5’-ATCCCTTATAAACGCTAACTGGAG-3’ + 5’-TCAAGTTGGATCTTCGGTTTATG-3’; P5: 5’-TACACCATGTCTTTTCCAGGAG-3’ + 5’-GATATAGAAGCTGAAGGAAGTTGATCT-3’; P6: 5’-AAAGATCCGGAGTTACACACTAGC-3’ + 5’-CATTCTGGTTCTTCACCAGTTTCT-3’. Sequences of oligonucleotides used for organellar DNA content quantification were previously described by Kim et al^41^.

### Phylogenetic analysis

Sequences of putative plant RNH1 proteins were retrieved from Phytozome^42^, CoGE^43^, Ensembl Plants^44^, PLAZA^45^ and 1KP^46^ through BLAST search and aligned in MAFFT^47^. Only sequences containing full length hybrid binding domain (HBD), RNASEH1-like domain and four amino acids (DEDD) crucial for catalytic activity^17^, were kept. BAli-Phy^48^ with default settings on CIPRES^49^ platform was used for phylogenetic analysis. Consensus tree was generated with 25% burn-in and visualized using FigTree. Subcellular localization was predicted by TargetP^26^.

### *In vitro* activity of RNH1s

6xHis-tagged HBDs and catalytic domains of RNH1A and B were expressed in *E. coli* from pET28 plasmid. Tagged proteins were bound on HisTrap (HP)(GE Healthcare) and eluted with increasing imidazole concentration. Enzymatic reaction was performed in Reaction Buffer (60mM NaCl, 16mM HEPES-KOH pH=7.0, 4% glycerol, 0.8mM DTT, 80µg/ml BSA, 4mM MgCl_2_) with 20nM fluor-labelled substrate and 2nM-2µM purified RNH1 in 20µl reaction. Reaction was conducted for 30min at 37°C and stopped by addition of EDTA to final concentration of 40mM. Products were analyzed on 15% denaturing TBE-urea polyacrylamide gels and visualized by fluorescence readout.

### Chlorophyll content measurement

Chlorophyll content was determined as described by Lichtenthaler^50^. Briefly, 1g of frozen 2-3 week old seedlings was ground in liquid nitrogen and chlorophyll was extracted with 100% acetone. Samples were centrifuged at 10000 x g for 10min at 4°C and absorbance was measured at 645 and 662nm. Chlorophyll concentration was calculated using following formula: C_a+b_(µg/ml)= 18.09 x A645 + 7.05 x A662.

### Microscopic observations

Mature siliques of Col-0 and *rnh1b*^*-/-*^ */ rnh1c*^*+/-*^ plants were destained in 70% ethanol and photographed under preparative microscope. For embryos observations siliques of Col-0 and *rnh1b*^*-/-*^ */ rnh1c*^*+/-*^ were dissected, seeds were split based on morphology and cleared for 2h at room temperature with Visikol^51^. Embryos inside seeds were visualized using Nomarski optics.

### Organellar DNA content quantification and expression analysis

Total DNA was isolated from 2-3 weeks old seedlings with DNeasy Plant Mini Kit (QIAGEN). 1ng of DNA per qPCR reaction was used as template. For expression analysis total RNA was extracted with RNeasy Plant Mini Kit (QIAGEN) from 2-3 weeks old seedlings and 500ng was treated with DNaseI (Ambion) and used for reverse transcription. 0.5ng of cDNA was used as template in qPCR. Significance was determined using confidence intervals.

### Replicative stress assay

Seeds were germinated on ½ MS plates supplemented with 1% sucrose and 0.75% agar. After 1 week seedlings were transferred to liquid ½ MS + 1% sucrose with 0 or 2mM hydroxyurea (HU) and grown in continuous light. After 2 weeks root length was measured.

## ACKNOWLEDGEMENTS

We want to thank Steven Smith for his help with phylogenetic analysis as well as Karolina Górecka and Małgorzata Figiel for their help with protein purification and biochemical activity assays. This work was supported by the National Science Foundation grant MCB 1120271 and the National Science Centre, Poland grant Polonez 2015/19/P/NZ1/03619 to A.T.W. This project has received funding from the European Union’s Horizon 2020 research and innovation programme under the Marie Skłodowska-Curie grant agreement No 665778.

## SUPPLEMENTAL DATA

**Figure S1.**
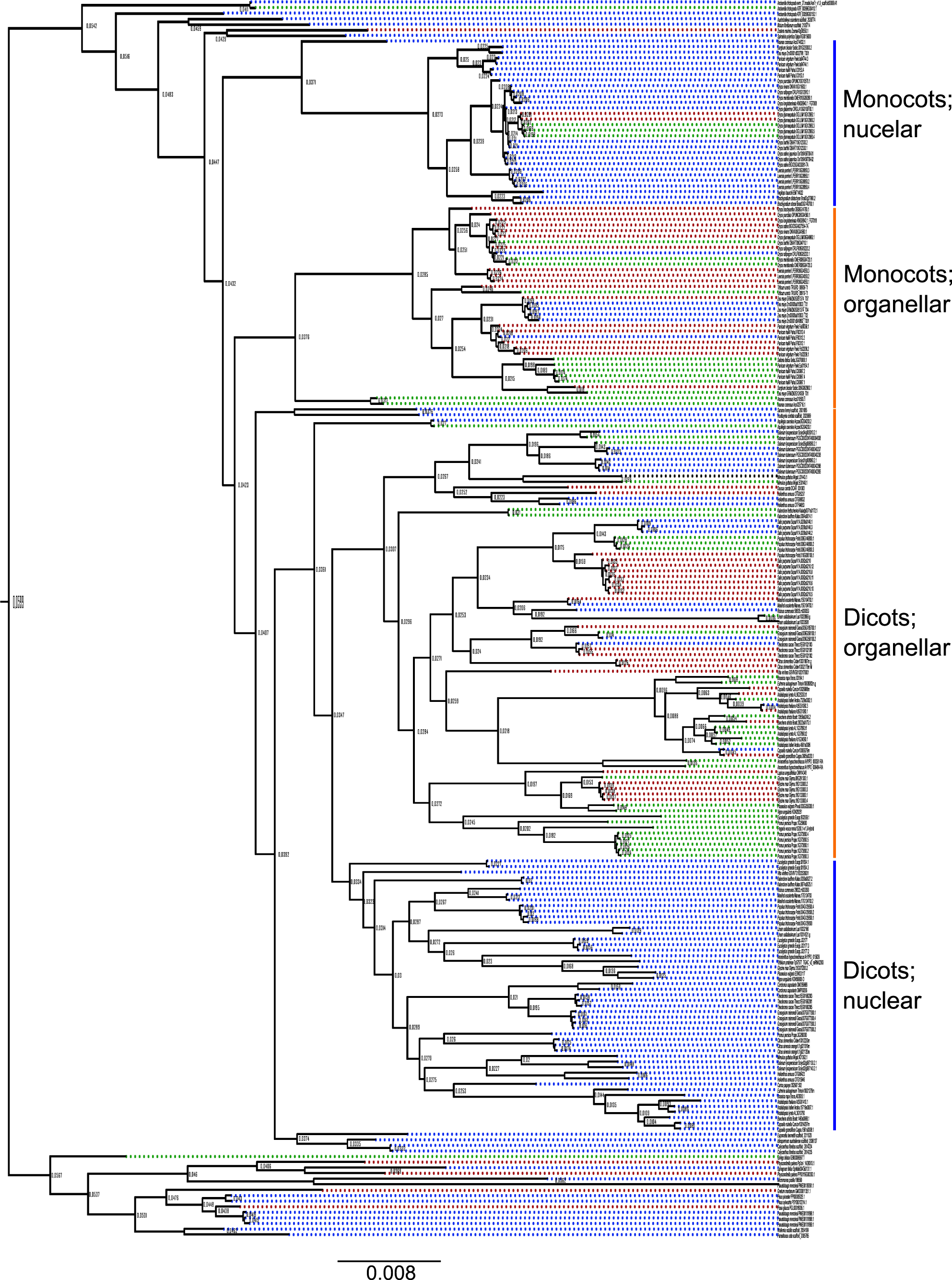
Evolutionary origin of plant RNase H1 proteins. Phylogenetic tree of all full length predicted RNase H1 proteins as shown in Fig. 1 with all species names and support values included.

**File S1**. Aminoacid sequences of all RNase H1 proteins shown in Fig. 1 and Fig. S1.

